# Transcriptomic Changes in Oligodendrocyte Lineage Cells During the Juvenile to Adult Transition in the Mouse Corpus Callosum

**DOI:** 10.1101/2024.03.28.587148

**Authors:** Tomonori Hoshino, Hajime Takase, Gen Hamanaka, Shintaro Kimura, Norito Fukuda, Emiri T Mandeville, Josephine Lok, Eng H Lo, Ken Arai

## Abstract

The corpus callosum, a major white matter tract in the brain, undergoes age-related functional changes. To extend our investigation of age-related gene expression dynamics in the mouse corpus callosum, we compared RNA-seq data from 2-week-old and 12-week-old wild-type C57BL/6J mice and identified the differentially expressed genes (e.g., *Serpinb1a, Ndrg1, Dnmt3a*, etc.) between these ages. Interestingly, we found that genes highly expressed in myelinating oligodendrocytes were upregulated in 12-week-old mice compared to 2-week-old mice, while genes highly expressed in oligodendrocyte precursor cells (OPCs) and newly formed oligodendrocytes were downregulated. Furthermore, by comparing these genes with the datasets from 20-week-old and 96-week-old mice, we identified novel sets of genes with age-dependent variations in the corpus callosum. These gene expression changes potentially affect key biological pathways and may be closely linked to age-related neurological disorders, including dementia and stroke. Therefore, our results provide an additional dataset to explore age-dependent gene expression dynamics of oligodendrocyte lineage cells in the corpus callosum.

## Introduction

The corpus callosum, the largest white matter structure in the brain, plays an important role in the inter-hemispheric transfer of sensory, motor, and cognitive information in most mammals ^1^. The corpus callosum is enriched in oligodendrocyte precursor cells (OPCs) and oligodendrocytes, and this region is implicated in various neurological disorders, such as Alzheimer’s disease ^2–5^. Throughout its lifespan, the corpus callosum undergoes significant changes. Studies have revealed intricate processes governing its formation, primarily through research on mice ^6–8^. In mice, myelination is observed starting from postnatal day 11, with rapid myelin formation occurring between days 14 and 45 ^9^. Furthermore, diffusion tensor imaging and computational neuroanatomical analyses reveal that fiber maturation (i.e., dense packing of myelinated axons) in the corpus callosum of C57BL/6J mice reaches a steady state between postnatal days 30 and 40 ^10^. In addition, the corpus callosum is a vulnerable region during aging ^11,12^.

White matter volume and myelin formation in the corpus callosum increase until adulthood, peaking before gradually decreasing with age ^13,14^. Studies using aged mice have shown activation of microglia and astrocytes ^15,16^, and demyelination has been found to activate microglia, thereby promoting the accumulation of amyloid-beta, a key factor in Alzheimer’s disease pathology ^17^. A systematic behavioral analysis using 3-to 22-month-old C57BL/6J mice revealed that the physical functions, such as locomotor activity, gait velocity, and grip strength, progressively decline, starting as early as 6 months of age in mice, while cognitive function begins to decline later, with considerable impairment present at 22 months of age ^18^. We have previously used 2-month-old and 8-month-old vascular dementia model mice (bilateral carotid artery stenosis [BCAS] mice) and found that middle-aged mice have more white matter damage after cerebral hypoperfusion by the BCAS surgery ^19^, suggesting that aging may contribute to greater vulnerability to white matter damage in the corpus callosum.

Although comprehensive analyses of the corpus callosum exist, these studies have important caveats. For instance, proteomic research has focused on early postnatal development within the first month after birth ^20^. Transcriptomic analyses have provided insights into the corpus callosum at various life stages, particularly in aging, as evidenced by studies on older mice ^15,16^. However, these studies tend to overlook the crucial transitional periods, such as the changes occurring between 2 weeks and 12 weeks postnatal. This oversight creates a significant gap in our understanding of the changes in the corpus callosum from postnatal to adult stages. The lack of detailed data across these critical transitions may hamper our ability to fully understand the biological foundations of age-related neurological disorders. While research has thoroughly investigated the development of the corpus callosum and its deterioration in later life, the intermediate stages remain understudied.

In this study, we aim to fill this gap by analyzing RNA-seq data from the corpus callosum of male C57BL/6J mice, a widely-used inbred strain, at 2 weeks of age and comparing it to the data at 12 weeks of age. In addition, we extended our analysis by comparing these findings with existing data sets from 5-week-old and 96-week-old mice. By examining the patterns of gene expression changes at these life stages, we seek to provide new insights into the aging process of the corpus callosum and its implications for neurological disorders.

## Results

### Isolation of corpus callosum from juvenile and young adult mice and the assessment

The diagram in Figure 1a summarizes the experimental approach in this study, i.e., expression levels of mRNA in the corpus callosum of juvenile (2-week-old) and young adult (12-week-old) mice were compared with RNA-seq experiments (Figure 1a). PCA analysis revealed that these samples were clustered into two groups (Figure 1b). To validate the successful isolation of the corpus callosum, we assessed the gene expression levels of specific markers for oligodendrocytes (*Mbp* and *Mobp*) and cortical neurons (*Reln, Rasfrf2, Pou3f2*, and *Foxp2*) and confirmed that the sample was white matter tissue based on the high expression of oligodendrocyte markers (Figure 1c).

**Figure 1.**
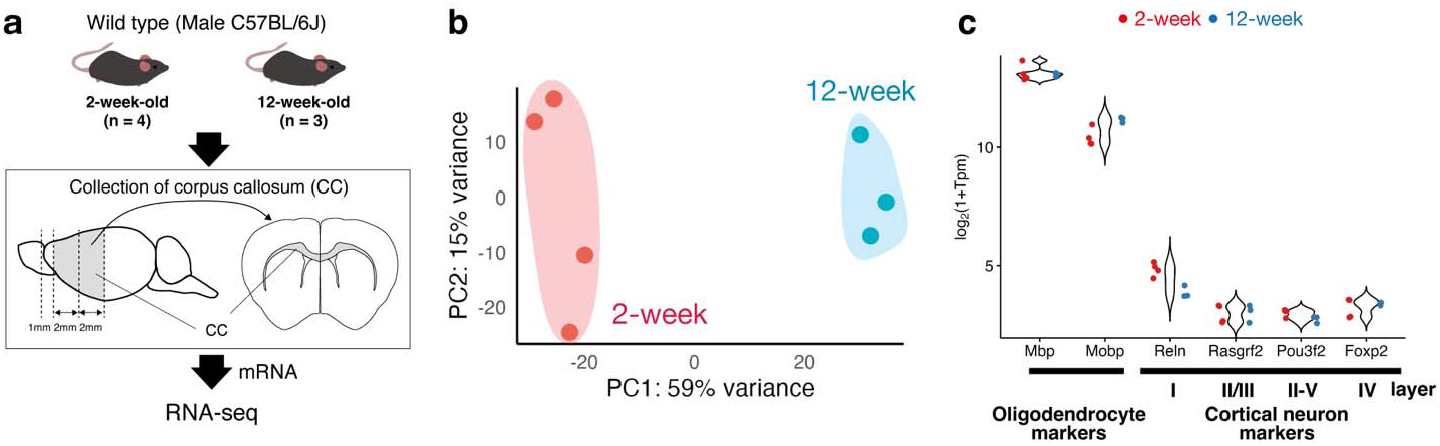
Corpus callosum sampling and RNA-seq analysis from juvenile to adult: (a) Schematic representation illustrating the collection of the corpus callosum (CC) in 2-week-old and 12-week-old mice. The isolated RNA was analyzed using RNA-seq. (b) Principal Component Analysis (PCA) plot of the samples. (c) Violin plots representing the expression of oligodendrocyte markers (*Mbp* and *Mobp*) and cortical neuron markers (*Reln* for layer I, *Rasgrf2* for layer II/III, *Pou3f2* for layer II-V, and *Foxp2* for layer IV).

### Transcriptome profiling of corpus callosum at different ages

We next explored the differentially expressed genes (DEGs), visualized in volcano plots (2-week vs. 12-week), and found that 439 genes were upregulated and 620 genes were downregulated in 12-week-old mice (a total of 1,059 genes; Figure 2a). The top 10 up-regulated genes by padj were *Serpinb1a, Gm21984, C030029H02Rik, Adamtsl4, Il33, Neat1, Ndrg1, Cbx7, Efhd1*, and *Qdpr*, while the top 10 down-regulated genes by padj were *Gng4, Col4a1, Dnmt3a, Bdh1, Apcdd1, Cd93, Nrep, Nid1, Emid1*, and *Marcksl1* (Figure 2b and 2c, Supplementary Table S1). When analyzing genes with notable expression in central nervous system cells (neuron, astrocyte, microglia, OPCs, newly formed oligodendrocytes, and myelinating oligodendrocytes) based on public RNA-seq datasets ^21,22^, the majority of upregulated DEGs were derived from myelinating oligodendrocytes, while the majority of downregulated DEGs were derived from OPCs and newly formed oligodendrocytes in 2-week vs. 12-week (Figure 2d).

**Figure 2.**
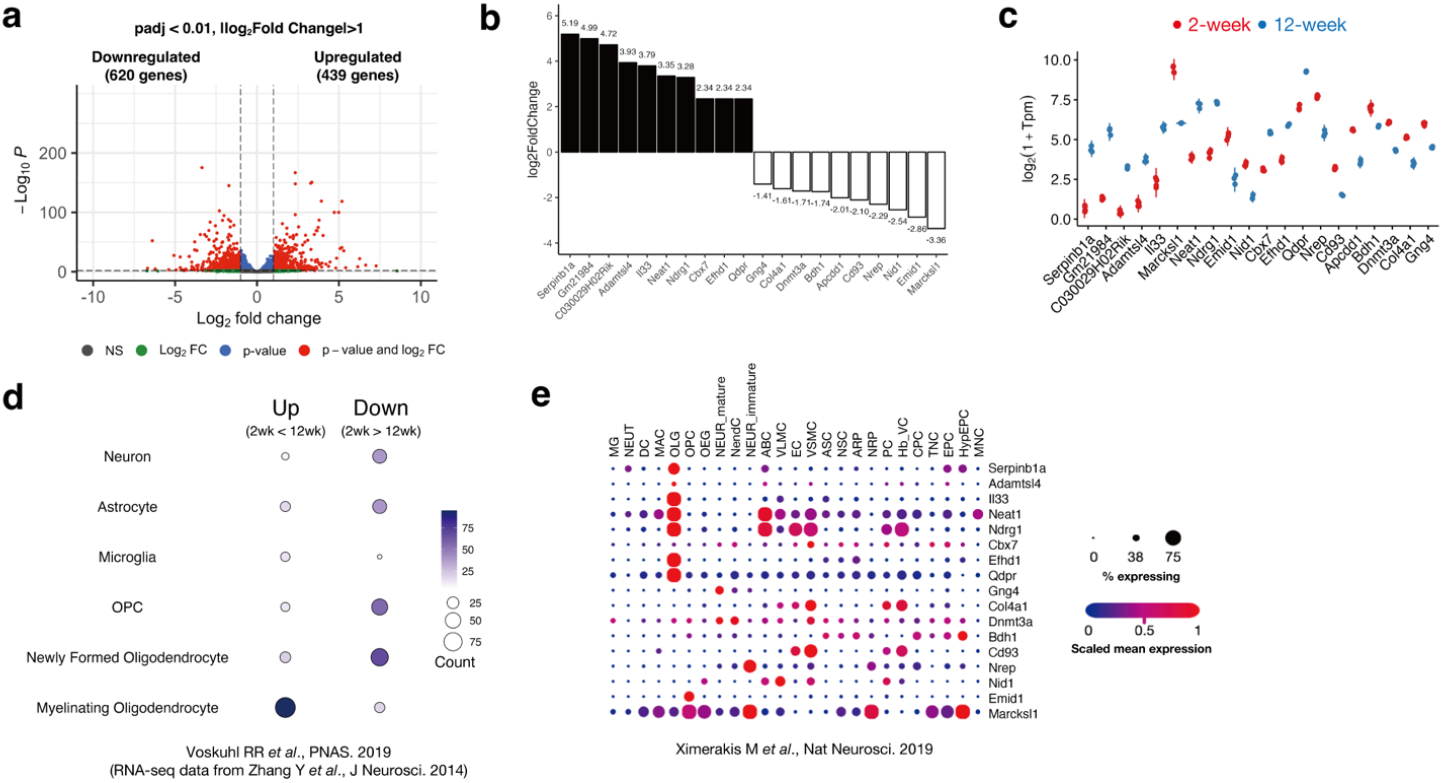
Transcriptome profiling of the corpus callosum from juvenile to adult: (a) Volcano plot illustrating DEGs (a total of 1,564 genes) between 2-week-old (n = 4) and 12-week-old (n = 3) mice. The DEGs cutoff was set at padj < 0.01 and | Fold Change (FC) | > 2 (|log2FC| > 1). (b) Bar plot of the top 10 genes upregulated and downregulated in 12-week-old WT mice compared to 2-week-old WT mice (by padj). (c) Violin plot of the top 10 genes upregulated and downregulated in 12-week-old WT mice compared to 2-week-old WT mice (by padj). (d) Classification of DEGs in 2-week vs. 12-week using cellular markers from previously published RNA-seq data ^21,22^. (e) Classification of the top 10 gene lists (*Gm21984, C030029H02Rik*, and *Apccd1* were not registered in this database) using public single-cell RNA-seq data ^23^. For abbreviations in this figure, see Supplementary Figure S2.

To further investigate the cell types of these highly up- or down-regulated genes, we used publicly available single-cell RNA-seq data from adult mouse brains (GSE129788) ^23^. Among the upregulated genes at 12 weeks of age, we verified that *Serpinb1a, Adamtsl4, Il3, Efhd1*, and *Qdpr* are predominantly expressed in oligodendrocyte cells, and *Ndrg1* and *Neat1* are in oligodendrocyte and vascular cells (ex. vascular smooth muscle cells [VSMC], endothelial cells [EC], and pericytes [PC]) (Figure 2e). On the other hand, among the downregulated genes, *Emid1* is predominantly expressed in OPC, *Nid1, Cbx7, Cd93*, and *Col4a1* are in vascular cells (ex. VSMC, EC, and PC), *Nrep* is in immature neurons, and *Gng4* are in mature neurons (Figure 2e).

## Gene ontology enrichment analysis of developmental-specific gene sets

**Gene** ontology (GO) analysis of the DEGs further revealed significant enrichment in various biological processes and pathways. Specifically, upregulated genes were significantly enriched in processes such as actin filament-based process, actin cytoskeleton organization, reversible hydration of carbon dioxide, actin filament organization, nitrogen metabolism in Mus musculus (house mouse), supramolecular fiber organization, regulation of interleukin-4 production, positive regulation of transcription in response to endoplasmic reticulum stress, protein refolding, and positive regulation of inflammatory response (Figure 3a, Supplementary Table S2). Downregulated genes were predominantly involved in extracellular matrix organization, external encapsulating structure organization, extracellular structure organization, extracellular matrix organization, ECM-receptor interaction in mus musculus (house mouse), mitotic cell cycle, mitotic cell cycle process, collagen formation, focal adhesion in mus musculus, and supramolecular fiber organization (Figure 3a, Supplementary Table S2).

**Figure 3.**
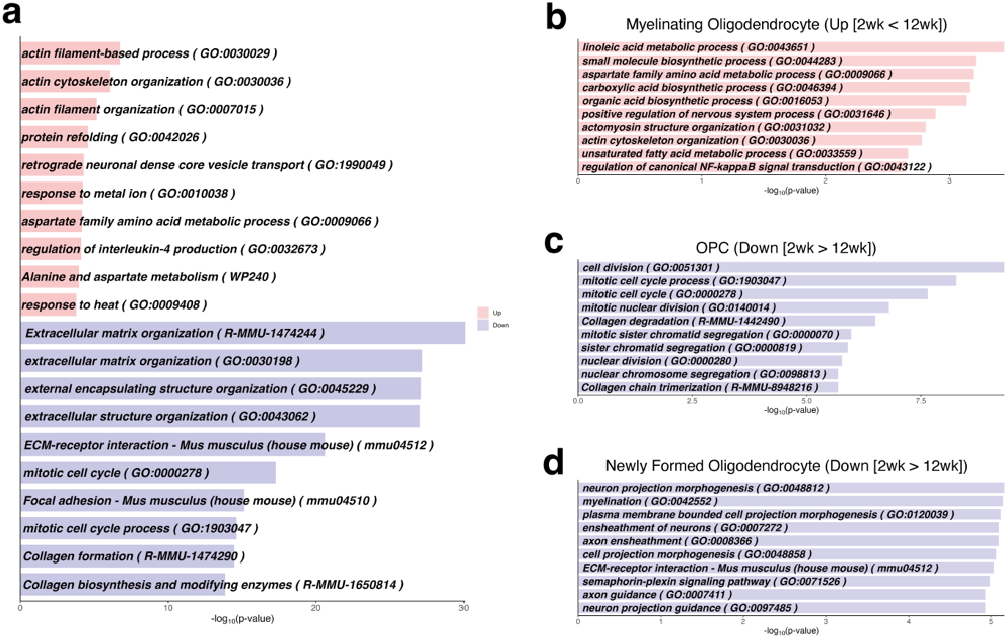
Gene ontology analysis of the corpus callosum from juvenile to adult: (a) Top 10 GO terms enriched among all DEGs in 2-week-old vs. 12-week-old. Upregulated genes (Up); red, Downregulated genes (Down); blue. (b) Top GO terms enriched among the some upregulated DEGs in 2-week-old vs. 12-week-old, which predominantly expressed in myelinating oligodendrocytes. (c) Top GO terms enriched among the some downregulated DEGs in 2-week-old vs. 12-week-old, which predominantly expressed in OPCs. (d) Top GO terms enriched among the some downregulated DEGs in 2-week-old vs. 12-week-old, which predominantly expressed in newly formed oligodendrocytes.

In Figure 2d, our analysis showed that among the genes with increased expression, 95 genes were abundantly expressed in myelinating oligodendrocytes (Supplementary Table S3). Additionally, among the genes with decreased expression, 56 genes were most abundantly expressed in OPCs, and 68 genes in newly formed oligodendrocytes (Supplementary Table S3). These subsets of genes were extracted and subjected to further GO analysis to elucidate the biological processes enriched in these specific cell populations. The GO analysis revealed that for myelinating oligodendrocytes, enriched biological processes included linoleic acid metabolic process, small molecule biosynthetic process, aspartate family amino acid metabolic process, carboxylic acid biosynthetic process, organic acid biosynthetic process, positive regulation of nervous system process, actomyosin structure organization, actin cytoskeleton organization, unsaturated fatty acid metabolic process, and regulation of canonical NF-kappaB signal transduction (Figure 3b, Supplementary Table S4). Among the genes downregulated in OPCs, processes such as cell division, mitotic cell cycle process, mitotic cell cycle, mitotic nuclear division, collagen degradation, mitotic sister chromatid segregation, sister chromatid segregation, nuclear division, nuclear chromosome segregation, collagen chain trimerization were prominent (Figure 3c, Supplementary Table S5). In newly formed oligodendrocytes, neuron projection morphogenesis, myelination, plasma membrane bounded cell projection morphogenesis, ensheathment of neurons, axon ensheathment, cell projection morphogenesis, ECM-receptor interaction - mus musculus (house mouse), semaphorin-plexin signaling pathway, axon guidance, and neuron projection guidance were among the top enriched processes (Figure 3d, Supplementary Table S6).

When we performed Gene Set Enrichment Analysis (GSEA) using a gene list ranked by fold change, we found that many functions related to the vascular system, such as circulatory system process, endothelial cell proliferation, vascular transport, endothelial cell migration, positive regulation of vasculature development, negative regulation of vasculature development, sprouting angiogenesis, regulation of vasculature development, vascular process in circulatory system, aorta development, artery development, and artery morphogenesis, were enriched among the genes that decreased from 2 weeks to 12 weeks of age (Supplementary Table S7, Supplementary Figure S1).

### Comparative analysis of corpus callosum DEGs from juvenile to adult and adult to aged mice

We also compared these DEGs (i.e., 2-between 2-week-old vs. 12-week-old) with our previous data sets of 20-week-old and 96-week-old mice (PRJNA1011381). The Venn diagram revealed that 13 genes are shared between these two comparisons (Figure 4a-d, Supplementary Figure S2). Among the common genes, 10 genes (*Spp1, Etnppl, C4b, Lyz2, Mki67, Tnc, Chst3, Gpr17, Kif19a*, and *Marcksl1*) showed consistent trends of either up or down-regulation from 2-to 12-week-old and from 20-to 96-week-old (Figure 4a-d). Furthermore, 1,046 genes are unique in the 2-week vs. 12-week comparison and 46 genes are unique in 20-week and 96-week comparison (Figure 4a). For example, among the top 10 DEGs of

**Figure 4.**
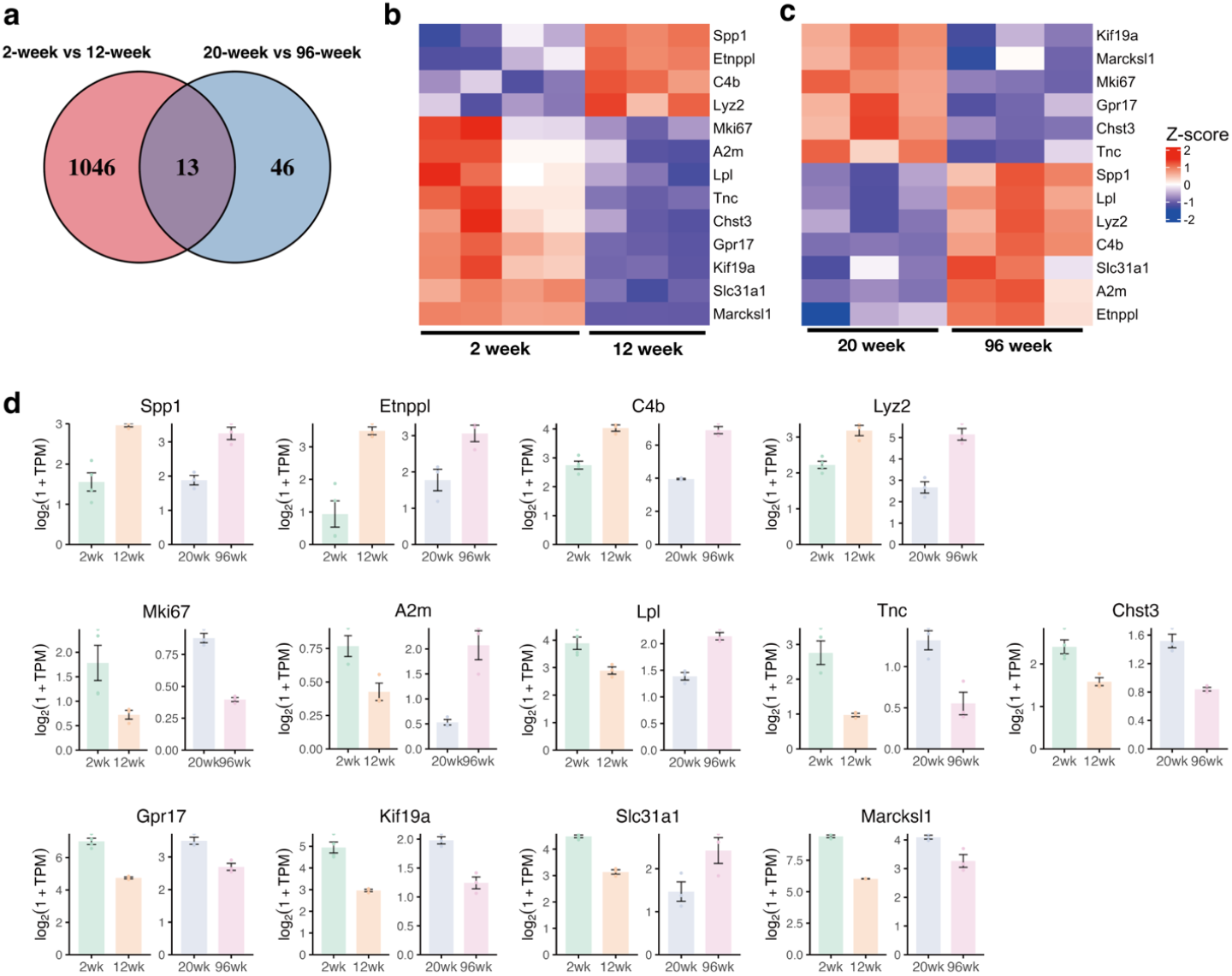
Comparative analysis of corpus callosum DEGs from juvenile to adult and adult to aged mice: (a) Venn diagram showing the overlapping DEGs between the comparisons of 2-week-old vs. 12-week-old and 20-week-old vs. 96-week-old. (b) Heatmap displaying the expression profiles of genes common between the DEGs of 2-week-old vs. 12-week-old and 20-week-old vs. 96-week-old, using data from 2-week-old and 12-week-old mice. (c) Heatmap displaying the expression profiles of genes common between the DEGs of 2-week-old vs. 12-week-old and 20-week-old vs. 96-week-old, using data from 20-week-old and 96-week-old mice. (d) Bar plot (with dot plot) of the common genes, using data from 2-week-old (2wk), 12-week-old (12wk), 20-week-old (20wk), and 96-week-old (96wk) mice. Error bars indicate SEM.

2-week vs. 12-week mice (10 upregulated and 10 downregulated in 12-wek mice, a total of 20 DEGs) (Figure 2b and 2c), 19 genes (*Serpinb1a, Gm21984, C030029H02Rik, Adamtsl4, Il33, Neat1, Ndrg1, Cbx7, Efhd1, Qdpr, Gng4, Col4a1, Dnmt3a, Bdh1, Apcdd1, Cd93, Nrep, Nid1*, and *Emid1*), excluding *Marcksl1*, were identified as DEGs unique to the 2- and 12-week comparison (Supplementary Figure S3a). Conversely, genes such as *Gfap, Col11a1, Cd68, B2m, Fcgr3, Podhb14, Cd84*, and *Mpeg1* were identified as DEGs unique to the 20-week vs. 96-week comparison (Supplementary Figure S3b).

## Discussion

In this study, we examined the expression profiles in the corpus callosum of mice and how these profiles change from juvenile to adult. While the age of mice cannot be directly translated to human age, a 0.5-month-old (∼2-week-old) mouse is roughly equivalent to a 0-1-year-old human, and a 3-month-old (∼12-week-old) mouse is roughly equivalent to a 15-20-year-old human ^24^. Notably, vascular dysregulation is observed at an early stage in patients with late-onset Alzheimer’s disease ^25^; therefore, understanding function at different ages is essential to elucidate the pathogenic mechanisms underlying these disorders. There have been analyses of expression variation during the postnatal to juvenile and adult to senescent periods ^15,16,20^; however, these studies may overlook the transitional periods between 2 and 12 weeks in the corpus callosum, one of the largest white matter regions, which plays an important role in the interhemispheric transfer of sensory, motor, and cognitive information. Therefore, this study provides an additional transcriptome profiling dataset to study the aging process of the corpus callosum and its implications in neurological disorders.

We observed that the expression of genes highly expressed in myelinating oligodendrocytes increased from juveniles to young adults, while the expression of genes highly expressed in OPCs and newly formed oligodendrocytes decreased (Figure 2d). This result is consistent with the previous electron microscopic and imaging evidence of myelin formation during this period ^9,10^, and one possibility for this result may be that it reflects the progression of major neurodevelopmental processes in the corpus callosum. Furthermore, GO analysis of the DEGs provided a deeper insight into the biological processes and pathways that are significantly affected during the developmental transitions in the corpus callosum. The upregulated genes in the corpus callosum from juvenile to young adult stages were particularly enriched in processes related to actin filament-based processes, actin cytoskeleton organization, actin filament organization, and related processes. These findings suggest a potential role for cytoskeletal reorganization and related molecular pathways in the maturation and functional modulation of the corpus callosum during these developmental stages. On the other hand, the downregulated genes were predominantly associated with extracellular matrix organization, external encapsulating structure organization, extracellular structure organization, extracellular matrix organization, and collagen formation, which may reflect the structural and compositional changes that occur in the extracellular environment of the corpus callosum as it matures. In addition, the downregulation of genes involved in the mitotic cell cycle and the mitotic cell cycle process may indicate a reduction in cellular proliferation, consistent with the observed decrease in the expression of genes highly expressed in OPCs and newly formed oligodendrocytes. We also found a number of genes that were common to the 2- and 12-week-old and 20- and 96-week-old comparisons, including *Spp1, Etnppl, Lyz2*, C4b, *Mki67, Tnc, Chst3, Gpr17, Kif19a*, and *Marcksl1* (Figure 4b-d). *C4b* has been reported to increase in the corpus callosum of aged mice ^15,16^, suggesting an important role of *C4b* in aged mice. Interestingly, some of the genes we identified as DEGs from either 2 weeks to 12 weeks or 20 weeks to 96 weeks are predominantly expressed in oligodendrocyte lineage cells. Since the corpus callosum is enriched in these cell types, and white matter dysfunction due to oligodendrocyte/myelin damage is one of the main features of age-related diseases, future studies are warranted to investigate how these genes change after white matter pathology.

Although our study provides a novel dataset to understand the changes in the transcriptome profiles of the mouse corpus callosum from juvenile to young adult stages, there are some caveats and limitations that we need to consider for future studies. First, our study included only male animals, which introduces a potential sex bias. Second, the use of bulk samples of the corpus callosum in this study introduces the possibility that significant changes in gene expression in some cell types may have been missed. The use of single-cell RNA sequencing in future studies would be valuable to examine the gene expression profiles of different cell types within the corpus callosum, thereby improving our understanding of its transcriptomic landscape. Third, this study focused on the gene expression changes of the corpus callosum, and we did not examine the cerebral cortex and hippocampus, which play important roles in various CNS diseases such as stroke, Alzheimer’s disease, and vascular dementia. Therefore, future studies are needed to investigate whether the gene expression changes we observed in this study are region-specific.

This study provides an additional resource for understanding development and aging in the corpus callosum. Our data set may help elucidate the molecular mechanisms that regulate white matter structure during development, including the role of oligodendrocyte lineage cells. In addition, since white matter is known to be a brain region vulnerable to aging, our dataset may be useful for future studies on the biological basis of aging and related neurological disorders.

## Methods

### Animals

We purchased pregnant female C57BL/6J mice from The Jackson Laboratory and housed them in a specific pathogen-free under a 12-hour light/dark cycle. The mice had ad libitum access to food and water throughout the experiment. For this study, we prepared the 2-week-old male C57BL/6J mice (n = 4) for RNA-seq, while the samples for the 12-week-old mice were derived from our previously reported experiments ^26^. In compliance with the National Institutes of Health (NIH) ethical guidelines, all experimental protocols involving animals were approved by the Institutional Animal Care and Use Committee at Massachusetts General Hospital.

### Tissue sampling and RNA extraction

Collecting of the corpus callosum from the brains was as previously described ^27^. Briefly, after euthanizing the mice, saline perfusion was performed using a pre-chilled solution, followed by decapitation. Brains were cooled in pre-chilled Hanks’ Balanced Salt Solution (Gibco, Billings, MT, USA, #14025-095) for a duration of 1 min. Sample collection was conducted between zeitgeber time; ZT3 and ZT5. RNA isolation from the corpus callosum was carried out using QIAzol® (QIAGEN, Venlo, Netherlands, #79306), according to the manufacturer’s instructions. The resultant pellet was dissolved in RNase-free water. NanoDrop spectrophotometers were used to assess the RNA quality and purity.

### RNA-sequencing

RNA samples were processed for RNA-seq, managed by Genewiz, Inc. (South Plainfield, NJ, USA). The library was prepared using a polyA selection for single indexing, and RNA-seq was performed on the Illumina HiSeq4000 platform (paired-end; 2 × 150 bp). The data were processed by mapping raw FASTQ files with STAR (version: 2.7.10a; mm10), followed by transcript quantification via RSEM (version: 1.3.3). Bioinformatics analysis was conducted in R (version: 4.3.0) using DESeq2 (version: 1.40.1) for differential expression analysis (|log2fold change (log2FC) | > 1, adjusted p-value (padj) < 0.01, mean base > 50). For most of our analyses, we used the normalized count data. However, for plotting, transcripts per kilobase million (TPM) values were used. Metascape was used for gene ontology (GO) analysis ^28^. The sequence data (FASTQ files) have been deposited under the BioProject accession for 2-week-old (PRJNA1086558) and 12-week-old (PRJNA727284; sham operation)^26^ (Supplementary Table S8). The RNA samples from the 2-week-old and 12-week-old mice were processed in the same batch (i.e., library preparation and sequencing were conducted simultaneously for these samples). Additionally, datasets from young (20-week-old) and aged (96-week-old) mice (PRJNA1011381) were also analyzed with the same threshold criteria. To match the sampling times of 2- and 12-week-old mice, we only used the sample at 1 hour after the onset of the light cycle (i.e.. ZT1) for detecting DEGs in 20-week-old and 96-week-old mice (n = 3).

## Acknowledgments

We thank our colleagues at the Neuroprotection Research Laboratories, Departments of Radiology and Neurology, Massachusetts General Hospital, and Harvard Medical School.

## Author’s contribution

T.H., H.T., and K.A. conceptualized and designed the study methodology. H.T., G.H., S.K., N.F., S.G., and E.T.M. provided essential resources for the research. T.H. performed the bioinformatics analysis and contributed to the visualization. T.H. and K.A. wrote the manuscript. E.H.L. and K.A. were involved in funding acquisition. All authors have read and approved the final manuscript.

## Data availability statement

The RNA-seq data have been deposited in the public repository under the accession code listed in the Methods. The datasets used and/or analyzed during the current study are available from the corresponding author upon reasonable request.

## Additional information

### Financial disclosure

This work was supported by a fellowship from the Uehara Memorial Foundation (FY 2022 to T.H.) and funding from the NIH.

### Competing Interests Statement

The authors declare that they have no competing interests.

### Supplemental material

Supplemental material for this article is available online.

## Notes

### Competing Interest Statement

The authors have declared no competing interest.

### Summary of Updates

The title and abstract were updated to clarify; Figure 2 and 3 were revised.

