## Supplemental material for "Transcriptomic Changes in Oligodendrocyte Lineage Cells During the Juvenile to Adult Transition in the Mouse Corpus Callosum"

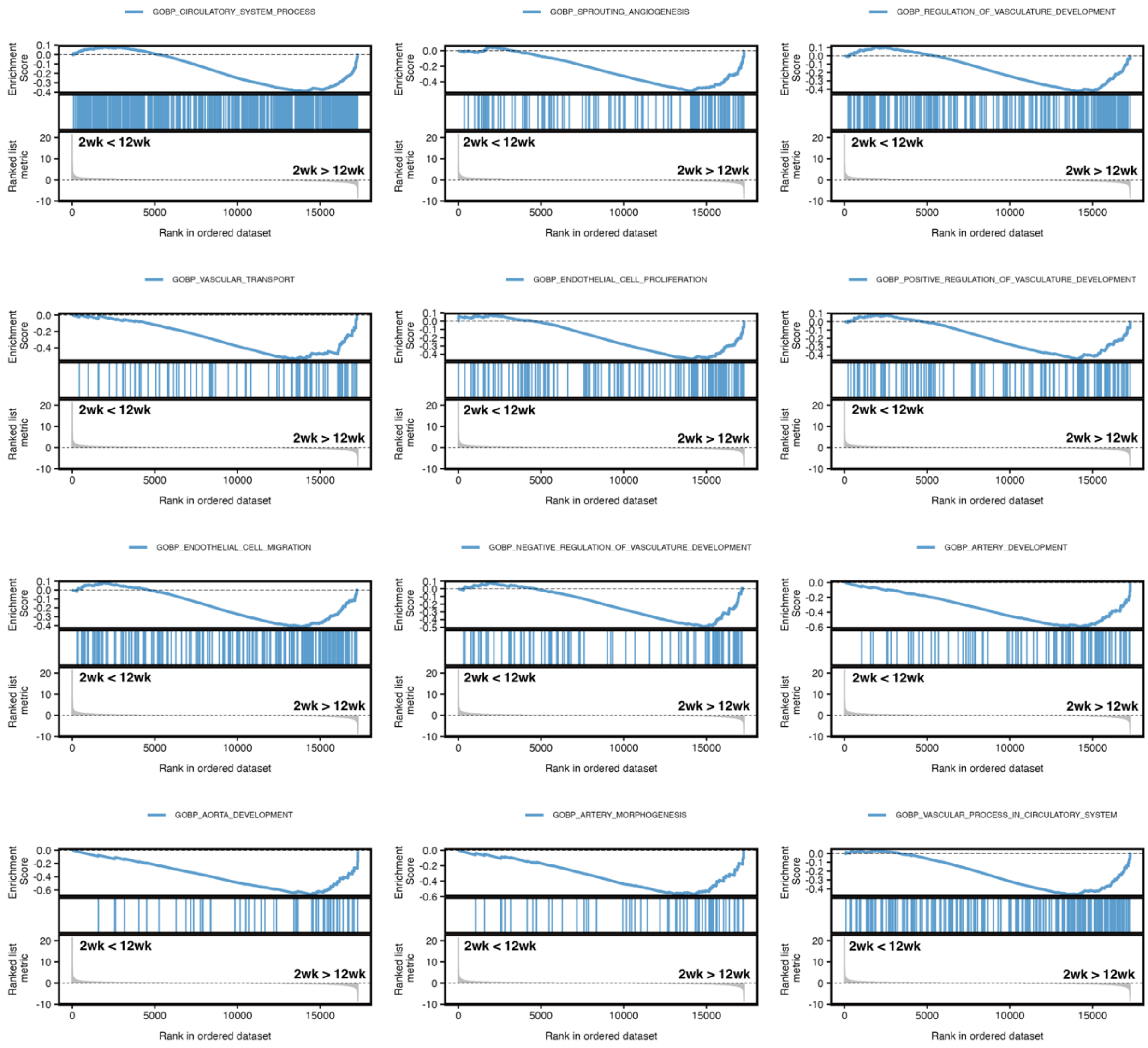


**Supplementary Figure S1. GSEA plot of corpus callosum from juvenile to adult mice:** GSEA plot showing enriched vascular-related functions among all ranked genes based on fold change from 2 to 12 weeks (Supplementary Table S8).

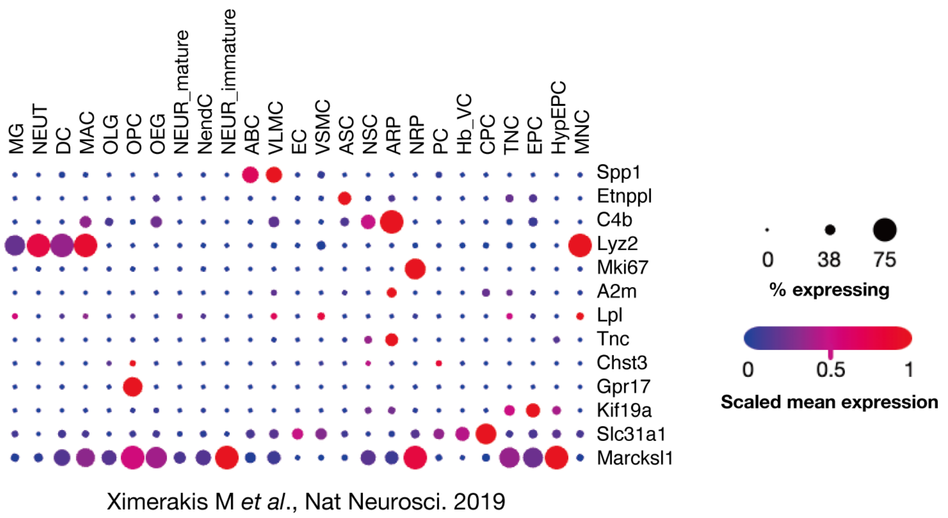


**Supplementary Figure S2. Classification of the common genes, using data from 2-week-old, 12-week-old, 20-week-old, and 96-week-old mice using public single-cell RNA-seq data [23].** The abbreviations used in this figure are as follows: In the oligodendrocyte lineage, OPC stands for oligodendrocyte precursor cells, OLG for oligodendrocytes, and OEG for olfactory ensheathing glia; in the astrocyte lineage, NSC represents neural stem cells, ARP for astrocyte-restricted precursors, and ASC for astrocytes; the neuronal lineage includes NRP as neuronal-restricted precursors, NEUR_immature for immature neurons, NEUR_mature for mature neurons, and NendC for neuroendocrine cells; ependymal cells have EPC as ependymocytes, HypEPC for hypendymal cells, TNC for tanycytes, and CPC for choroid plexus epithelial cells; vasculature cells comprise EC for endothelial cells, PC for pericytes, Hb-VC for hemoglobin-expressing vascular cells, VSMC for vascular smooth muscle cells, VLMC for vascular and leptomeningeal cells, and ABC for arachnoid barrier cells; and for immune cells, MG indicates microglia, MNC for monocytes, MAC for macrophages, DC for dendritic cells, and NEUT for neutrophils.

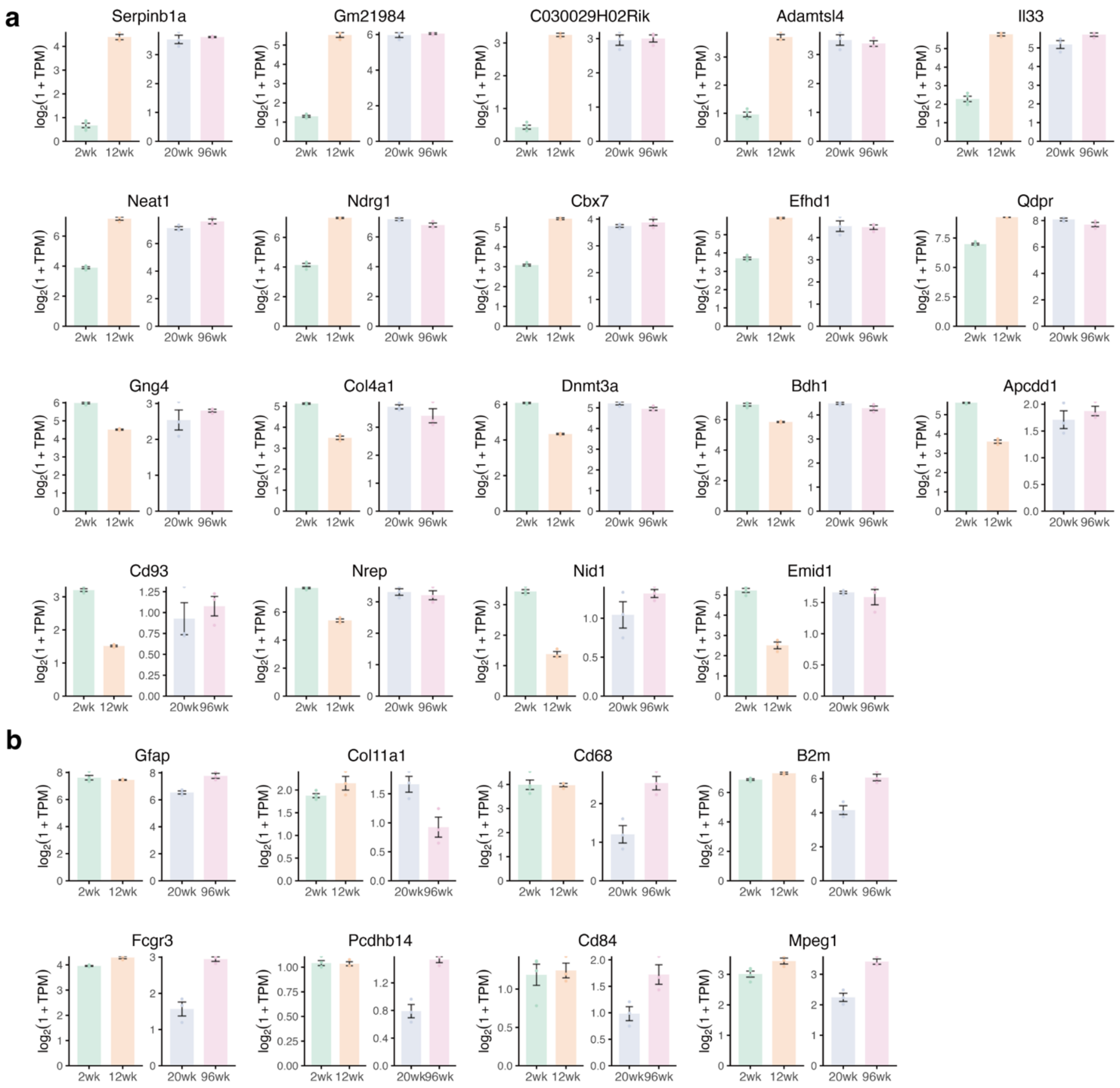


**Supplementary Figure S3. Analysis of unique DEGs in comparisons from juvenile to adult and adult to aged mice:** (**a**) Bar plots (with dot plots) showing the top 10 upregulated and downregulated genes unique to the 2-week vs. 12-week comparison, not detected in the 20-week vs. 96-week comparison, using data from 2-week-old (2wk), 12-week-old (12wk), 20-week-old (20wk), and 96-week-old (96wk) mice. (**b**) Bar plots (with dot plots) showing some genes detected as DEGs uniquely in the 20-week vs. 96-week comparison, using data from 2-week-old (2wk), 12-week-old (12wk), 20-week-old (20wk), and 96-week-old (96wk) mice. Error bars represent SEM.

**Supplementary Table:**

**Supplementary Table S1. Raw count data of the corpus callosum in 2-week-old and 12-week-old mice.**

**Supplementary Table S2. DESeq2 result of the corpus callosum in 2-week-old and 12-week-old mice.**

**Supplementary Table S3. Gene ontology analysis of the corpus callosum from juvenile to adult:** Top 10 GO terms enriched among all DEGs in 2-week-old vs. 12-week-old. Up: Upregulated genes. Down: downregulated genes.

**Supplementary Table S4. The list of DEGs enriched in myelinating oligodendrocytes, OPCs, and newly formed oligodendrocytes.**

**Supplementary Table S5. Top GO terms enriched among the some upregulated DEGs in 2-week-old vs. 12-week-old, which predominantly expressed in myelinating oligodendrocytes:** Up; Upregulated genes.

**Supplementary Table S6. Top GO terms enriched among the some downregulated DEGs in 2-week-old vs. 12-week-old, which predominantly expressed in OPCs:** Down; downregulated genes.

**Supplementary Table S7. Top GO terms enriched among the some downregulated DEGs in 2-week-old vs. 12-week-old, which predominantly expressed in newly formed oligodendrocytes:** Down; downregulated genes.

**Supplementary Table S8. GSEA results among all ranked genes based on fold change from 2 to 12 weeks.**
